# Single-Cell transcriptional changes in hypothalamic CRH-expressing neurons after early-life adversity inform enduring alterations in responses to stress

**DOI:** 10.1101/2021.08.31.458231

**Authors:** Annabel K Short, Christina Wilcox, Yuncai Chen, Aidan L Pham, Matthew T Birnie, Jessica L Bolton, Ali Mortazavi, Tallie Z. Baram

## Abstract

Mental and cognitive health, as well as vulnerability to neuropsychiatric disorders, involve the interplay of genes with the environment, particularly during sensitive developmental periods. Early-life stress / adversity (ELA) promotes vulnerabilities to stress-related affective disorders, yet it is unknown how a transient ELA dictates life-long neuroendocrine and behavioral reactions to stress. The population of hypothalamic corticotropin-releasing hormone (CRH)-expressing neurons that regulate stress-responses is a promising candidate to mediate the enduring influences of ELA on stress-related behavioral and hormonal responses via enduring transcriptional and epigenetic mechanisms. Capitalizing on a well-characterized model of ELA, we examined here the ELA-induced changes in gene expression profiles of stress-sensitive CRH-neurons in the hypothalamic paraventricular nucleus (PVN) of male mice. Given the known heterogeneity of these neuronal populations, we employed single-cell RNA sequencing (RNA-seq) approaches. The use of single-cell transcriptomics identified distinct CRH-expressing neuronal populations characterized by both their gene expression repertoire and their neurotransmitter profiles. Expression changes provoked by ELA clustered around genes involved in neuronal differentiation, synapse formation, altered energy metabolism and the cellular responses to stress and injury. Notably, the ELA-induced transcriptional changes took place primarily in subpopulations of glutamatergic CRH cells. Finally, ELA-induced transcriptional reprogramming of hypothalamic CRH-expressing neurons heralded significant, enduring disruptions of both hormonal and behavioral responses to stress throughout life.

## Introduction

Mental and cognitive health as well as vulnerability to neuropsychiatric disorders involve the interplay of genes with the environment during sensitive developmental periods (Bale et al. 2010; Klengel & Binder 2015; Short & Baram 2019). Genetic and environmental factors contribute to the development and maturation of neurons, synapses, and brain circuits, which, in turn drive long-lasting phenotypes.

Early-life adversity (ELA) promotes vulnerability to stress throughout life, but the mechanisms for these enduring phenotypes are incompletely understood (Lupien et al. 2009; Bale et al. 2010; Franklin et al. 2012; Short & Baram 2019). Stress-sensitive corticotropin-releasing hormone (CRH)-expressing neurons residing in the hypothalamus are candidate mediators of the long-lasting effects of ELA as they contribute to both behavioral and hormonal response to stress (Joëls & Baram 2009; Ulrich-Lai & Herman 2009; Füzesi et al. 2016; Kim et al. 2019; Daviu et al. 2020b). The release of CRH from the hypothalamic paraventricular nucleus (PVN) neurons induces pituitary adrenocorticotropic hormone (ACTH) secretion, which stimulates the release of corticosterone from the adrenal glands (Vale et al. 1981; De Kloet et al. 2005; McEwen 2007). CRH release may also regulate the stress response within the PVN by signaling to local CRH responsive neurons (Jiang et al. 2018; Jiang et al. 2019).

CRH expressing PVN neurons have traditionally been divided into three major subpopulations (Swanson et al. 1987; Tasker & Dudek 1991; Aguilera & Liu 2012; Biag et al. 2012). Pre-autonomic, non-neurosecretory PVN neurons project to the brainstem and spinal cord (Swanson & Kuypers 1980). Magnocellular neurons primarily secrete arginine vasopressin (AVP) or oxytocin into circulation and project to the posterior pituitary (Hoffman et al. 1991). The predominant population of CRH cells within the PVN are neuroendocrine (Wamsteeker Cusulin et al. 2013), and these neurosecretory parvocellular neuroendocrine cells project to the median eminence and release CRH potentially with other peptides (Swanson et al. 1987). In contrast to this traditional classification, recent single-cell analyses of hypothalamic transcriptomes have identified multiple molecular-defined clusters of neurons that express CRH within the PVN and throughout the hypothalamus that may represent different subpopulations or distinct neuronal functional states (Romanov et al. 2017a; Romanov et al. 2017b; Kim et al. 2020; Romanov et al. 2020; Xu et al. 2020).

Early-life experiences modulate CRH expression levels throughout the brain (Francis et al. 1999; Chen & Baram 2016). In the PVN specifically, optimal rearing conditions, including augmented maternal care, repress CRH expression (Fenoglio et al. 2004; Plotsky et al. 2005; Korosi et al. 2010), whereas adversity in early life may increase (Plotsky et al. 2005; Gunn et al. 2013) or decrease (Rice et al. 2008) the peptide’s expression. While the mechanisms by which environmental and maternal signals regulate CRH expression remain unclear, these signals influence the innervation of CRH-expressing cells. Within the PVN, augmented maternal signals reduce excitatory synapses to CRH cells (Korosi et al. 2010; Singh-Taylor et al. 2018), whereas adversity, including physical stress and unpredictable early-life environment, increases excitatory glutamatergic transmission onto the same cell population (Gunn et al. 2013; Bolton et al. 2021).

Synaptic neurotransmission induces signals to the nucleus and initiates transcriptional programming (West et al. 2002; Mohammad & Baylin 2010; Karsten & Baram 2013). Indeed, the epigenetic / transcriptional effects of early-life adversities on neurodevelopment have been well documented (Suderman et al. 2012; Hunter & McEwen 2013; Lucassen et al. 2013; Szyf 2013; Bale 2015; Deussing & Jakovcevski 2017; Peña et al. 2017; Gray et al. 2018; Bolton et al. 2020). ELA influences stress responses enduringly and these responses are governed by the stress-sensitive CRH cells in PVN (Avishai-Eliner et al. 2001; Fenoglio et al. 2006; van Bodegom et al. 2017; Babicola et al. 2021). However, it remains unclear how ELA influences gene expression programs in these neurons, if such changes are specific to sub-populations of CRH-expressing neurons, and whether the transcriptional changes are associated with increased vulnerabilities to stress throughout life.

Here we employed a model of ELA which provokes major alterations in cognitive and emotional outcomes (Gilles et al. 1996; Avishai-Eliner 2002; Brunson et al. 2005; Ivy et al. 2008; Ivy et al. 2010; Wang et al. 2011; Molet et al. 2014; Molet et al. 2016a; Bath et al. 2016; Heun-Johnson & Levitt 2016; Walker et al. 2017; Bolton et al. 2018a; Bolton et al. 2018b; Santiago et al. 2018; Short & Baram 2019; Goodwill et al. 2019; Short et al. 2020), including augmented responses to stress (Rice et al. 2008; Molet et al. 2016b; Bolton et al. 2021). We focused on the change in gene expression profiles of stress-sensitive PVN-CRH-neurons following ELA in male mice. We employed single-cell RNA sequencing (RNA-seq) to probe the effects of ELA on gene expression programs in distinct neuronal populations, determine the potential selectivity of the effects of ELA and identify their downstream consequences.

We found that CRH-expressing populations of neurons in the PVN cluster by both their gene expression profiles and their neurotransmitter phenotype. ELA significantly modified the gene expression profiles of these neurons, reducing programs of neuronal development and differentiation and enhancing gene families involved in responses to stress and inflammation. The use of single-cell transcriptomics revealed novel subpopulations of CRH cells and established the impact of ELA on gene expression profiles as cell-type specific, with distinctive influence on the different clusters and subpopulations of CRH neurons. Importantly, ELA-provoked transcriptional alterations are biologically impactful, as they translate into phenotypic changes, including augmented behavioral and neuroendocrine responses to stress later in life.

## Results

### CRH-expressing neurons in the developing mouse hypothalamus belong to distinct populations with unique gene-expression profiles

The hypothalamic PVN is known to harbor several types of CRH-expressing neurons, including a population of cells that directly release CRH into the portal bloodstream and others projecting to specific brain areas (Swanson et al. 1987; Aguilera & Liu 2012). Within the PVN, the majority of CRH-expressing (CRH+) cell populations co-express glutamate, a key excitatory neurotransmitter (Hrabovszky et al. 2005; Hrabovszky & Liposits 2008), whereas others express genes associated with inhibitory GABAergic neurotransmission (Meister et al. 1988). To delineate these distinct populations in developing mouse PVN and determine their potential contribution to the phenotypic consequences of ELA, we employed single-cell transcriptomics. As described in Figure 1, we reared mice in cages with limited bedding and nesting, a well-characterized model of ELA or in typical cages during the first week of life (Figure 1A). Individual cells from a tissue block containing the PVN (Figure 1B) were obtained from 10-to 12-day old mice reared in either type of environment. The cells were FACS sorted and sequenced individually (Figure 1B).

**Figure 1.**
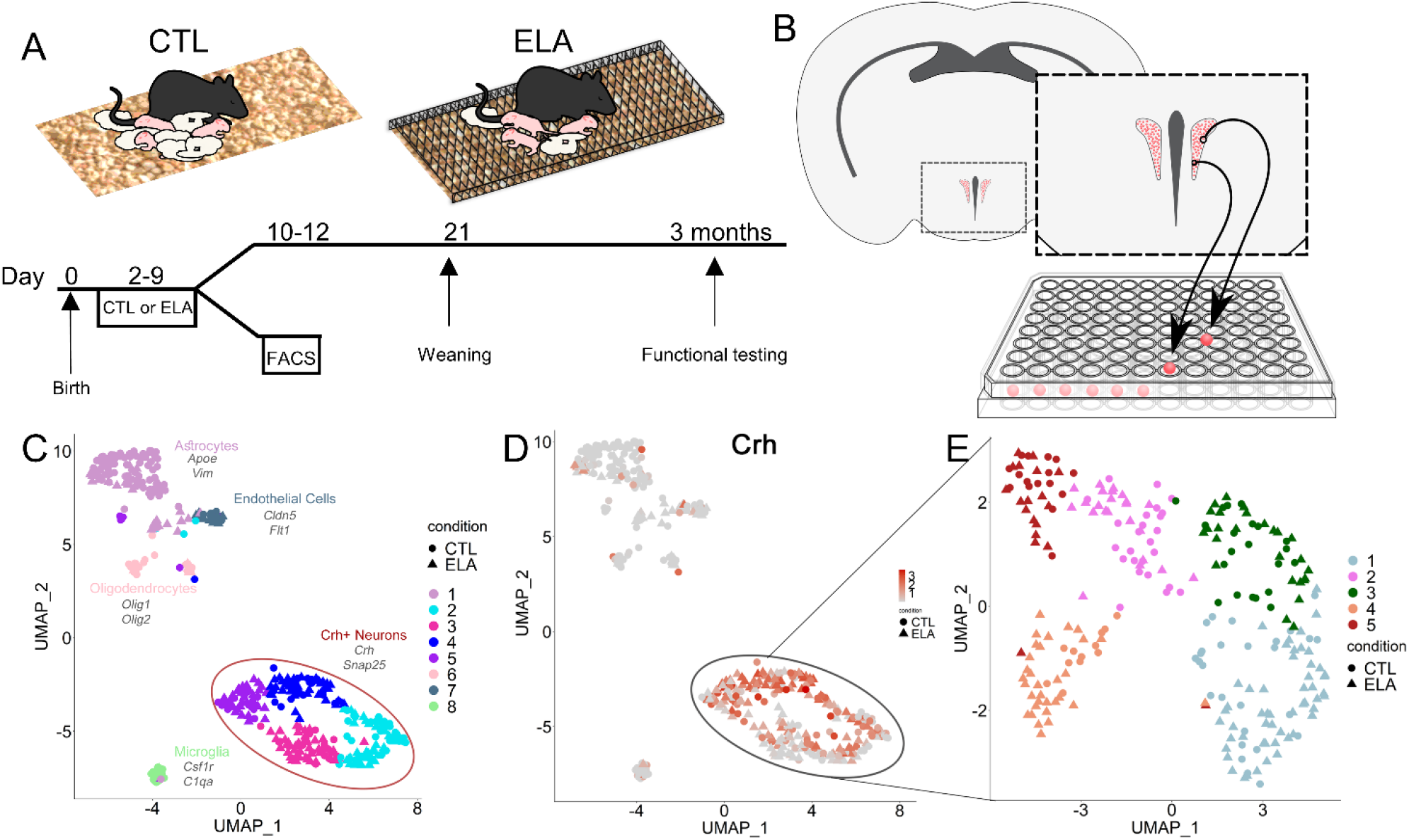
Single-cell RNA sequencing of CRH cells in the hypothalamic paraventricular nucleus reveals distinct neuronal populations. (A). Mice exposed to either control or early-life adversity conditions during the ten days of life were either kept for functional testing or employed for collection of PVN cells during postnatal days 10-12. (B). CRH cells were collected from Crh-IRES-Cre; Ai14 tdTomato mice by dissecting a PVN-containing tissue block, sorted into individual wells using FACS, and sequenced. Uniform Manifold Approximation and Projection (UMAP) was performed to cluster sorted cells and revealed 8 distinct groups of cells, the majority of which were classified as neurons based on expression of *Snap25* (C). (D). These neurons, the majority of CRH expressing cells, further clustered into 5 sub-populations.

We first ascertained that harvested cells did not cluster based the age or weight of the mice or on the harvest batch (Supplementary Fig 1A-C). Then, individual cells passing filter criteria were characterized as CRH-expressing neurons or non-neuronal cells including microglia, astrocytes and endothelial cells, based on their expression of marker genes (Figure 1C), and the latter were excluded from further analyses. We also excluded cells expressing *Pgr15l*, which is co-expressed with CRH in GABAergic neurons residing at the border of the dorsomedial hypothalamus and the PVN (Supplementary Figure 1D) (Romanov et al. 2017a; Romanov et al. 2017c). Focusing on cells co-expressing *Crh* and the neuronal synaptic protein *Snap25*, (Figure 1D) we characterized this neuronal population using share nearest neighbor clustering (using the Seurat package) (Stuart et al. 2019), and this analysis segregated the CRH+ neurons into five distinct clusters (Figure 1E).

### Early life adversity significantly impacts gene expression programs in hypothalamic CRH cells

The overall distribution of CRH+ cells within clusters did not distinguish between mice reared under control conditions and those experiencing a week of early-life adversity during a sensitive developmental period (*X*^2^_(4, *N*=254)_ =7.42, p=0.11) (Figure 2A); there was slight underrepresentation in cluster 2 and over-representation in cluster 4 of ELA cells compared to what would be expected by chance (Supplementary Fig 2A). In contrast, transcriptional analyses of ELA cells compared to CTL cells revealed profound changes in the expression of several key genes and gene families (Figure 2B-C). Using an FDR <0.1, 46 genes were differentially expressed in the overall CRH+ population of control and ELA mice. Of these, 28 were higher in the controls and 18 in the ELA (Figure 2B).

**Figure 2.**
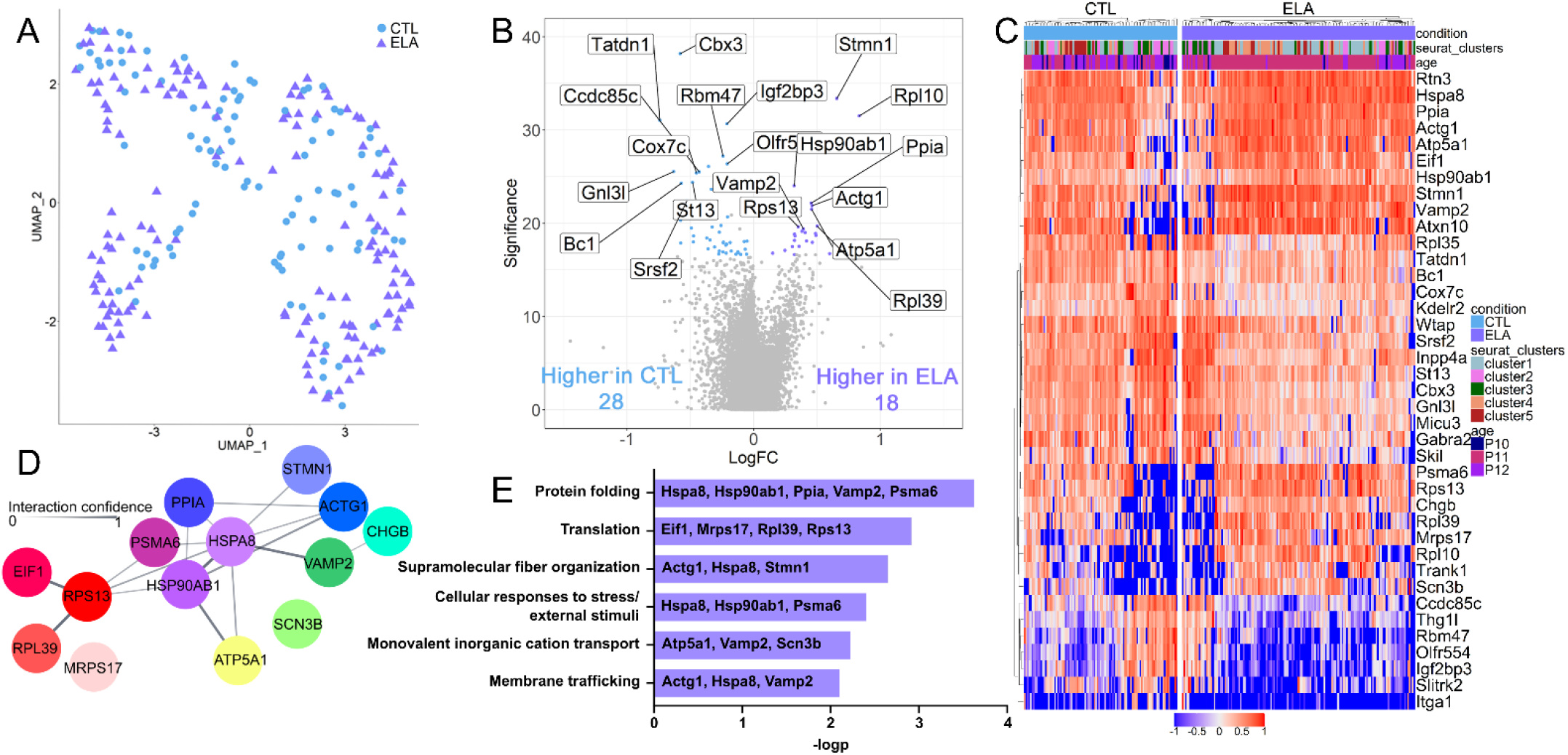
Transcriptomic changes induced by ELA in CRH neurons of the PVN. (A). Cells from CTL (102) and ELA (152) mice are similarly distributed across the UMAP clusters. (B). The volcano plot identifies genes that are significantly enriched (log fold change) in CTL (blue) and ELA (purple) cells. (C). ELA and CTL cells have different transcriptomic profiles that are independent of age or cluster (log fold change by row). (D). Predicted protein-protein interaction (cytoscape) of genes enriched in ELA. Nodes represent proteins and edges represent predicted interaction; line strength indicates confidence of predicted interactions between 0-1). Cool colors represent genes associated with neuron activation, and warm colors with translation and protein folding. (E). Metascape analysis revealed genes significantly (- logp) associated with predicted pathways as identified by gene ontology.

String network (Doncheva et al. 2019) and gene ontology enrichment (Zhou et al. 2019) analyses of genes differentially upregulated in ELA mice revealed groups of genes with predicted protein interactions (Figure 2D) and functional pathway associations (Figure 2E). Genes upregulated following ELA included pathways involved in cellular response to stress and in protein folding (Figure 2D-2E). Two heat-shock proteins *Hspa8*, Heat Shock Protein Family A Member 8; and *Hsp90ab1*, Heat Shock Protein 90 KDa, which are members of the heat shock protein 90 family associated with response to environmental stressors (Jarosz & Lindquist 2010) were augmented, as well as *Psma6* (Proteasome 20S Subunit Alpha 6) and *Ppia* (Peptidylprolyl Isomerase A). There was also enrichment of genes associated with regulation of synaptic vesicle content and transport (*Vamp2*, Vesicle Associated Membrane Protein 2; *Chgb*, Chromogranin B; *Atp5a1*, ATP Synthase F1 Subunit Alpha; and *Scn3b*, Sodium Voltage-Gated Channel Beta Subunit 3), membrane trafficking (*Atp5a1, Vamp2, Scn3b*) and neuronal structure (*Actg1*, Gamma-actin; *Hspa8*; and *Stmn1*, Stathmin 1) (Figure 2D-E). String analyses demonstrated predicted protein interactions of the upregulated heat-shock proteins and the products of genes associated with the multiple enriched pathways (Figure 2D). In cells from ELA mice, pathway analysis also confirmed upregulation of translation (*Eif1*, Eukaryotic Translation Initiation Factor 1; *Mrps12*, 28S ribosomal protein S12 mitochondrial; *Rpl39*, Ribosomal Protein L39; and *Rps13*, Ribosomal Protein S13) (Figure 2E). In contrast to the several gene networks enriched in ELA cells, as described above, network analyses of genes upregulated in CTL cells did not reveal significantly interacting genes or enriched pathways.

### Gene expression programs are differentially influenced by early-life adversity in distinct subpopulations of CRH expressing cells

CRH+ cells in the PVN belong to distinct populations with distinct functional outputs (Swanson et al. 1987; Aguilera & Liu 2012; Xu et al. 2020). Therefore, we examined if specific populations of the individually analyzed cells were differentially influenced by ELA. To better identify potential neuronal subpopulations, we first delineated the genes whose expression defined individual cell clusters (Figure 1E). Genes enriched in cluster 1 included *Avp* (Arginine Vasopressin), the gene encoding the stress-related neuropeptide vasopressin, as well as the stress-steroid receptors *Nr3c1* (mineralocorticoid receptor, MR) and *Nr3c2* (glucocorticoid receptor, GR), *Adra1b* (alpha-adrenergic receptor 1B), and ion channels/transporters including *Kcna5* (Kv1.5, Potassium Voltage-Gated Channel Subfamily A Member 5) and *Slc12a7* (KCl cotransporter 4, KCC4) (Supplementary Fig 2B). Cluster 2 was enriched with genes associated with GABAergic neurotransmission including *Gad1* (Glutamate Decarboxylase 1) and *Gad2* (Glutamate Decarboxylase 2) and also expressed *Nts* (Neurotensin) (Supplementary Fig 2B). Similar to cluster 1, cluster 3 cells had enriched expression of *Avp*, the steroid receptor *Nr4a1* (Nuclear Receptor Subfamily 4 Group A Member 1) as well as ion channels/transporters including *Scn7a* (Nav2.1, Sodium Voltage-Gated Channel Alpha Subunit 7) and *Slc4a11* (Bicarbonate Transporter Related Protein 1) (Supplementary Fig 2B). Cluster 3 was also characterized by high expression of the vasopressin secretion associated gene *Npff* (Neuropeptide FF-Amide Peptide Precursor) (Supplementary Fig 2B). Cells in cluster 4 had the most diverse expression profile including high expression of *Ntng1* (Netrin G1), markers of glutamatergic neurotransmission such as *Slc17a6* (vGLUT2) and *Gls* (Glutaminase), and the glutamate receptors *Grin3a* (Glutamate Ionotropic Receptor NMDA Type Subunit 3A, GluN3A) and *Grin2b* (Glutamate Ionotropic Receptor NMDA Type Subunit 3B, GluN3B), the GABA-A receptors *Gabra4* (Gamma-Aminobutyric Acid Type A Receptor Subunit Alpha4) *and Gabra5* (Gamma-Aminobutyric Acid Type A Receptor Subunit Alpha5), the cholinergic receptor *Chrna4* (Cholinergic Receptor Nicotinic Alpha 4 Subunit), and multiple ion channels/transporters (*Kcnd2*, Potassium Voltage-Gated Channel Subfamily D Member 2; *Clcn4*, Chloride Voltage-Gated Channel 4; *Slc29a1*, ENT; *Slc20a1*, Pit-1; *Slc8a1*, NCX1; *Slc12a5*, KCC2; *Slc25a22*, Mitochondrial Glutamate Carrier 1; and *Slc6a13*, GABA Transporter 2) (Supplementary Fig 2B). Cluster 4 also had high expression of *Maoa* (Monoamine Oxidase A). Cluster 5 had the fewest differentially expressed genes and was specifically characterized by the expression of the GABA transporter *Slc6a1* (GABA Transporter 1) (Supplementary Fig 2B).

The cluster analysis described above was based on CRH+ neurons from both ELA and CTL populations. Therefore, to exclude the possibility that ELA might modulate the expression state or clustering of the population of PVN-CRH cells in the developing mouse, we performed the same cluster analysis on CTL cells only. Given that the cell number was now lower, the results - three clusters that encompassed most genes from the above clusters (Supplementary Fig 3) - supported the validity of this clustering approach: CTL cluster 1 contained genes from clusters 1 and 3, CTL cluster 2 contained genes also expressed in cluster 2. The third CTL cluster contained more unique genes but also included genes from clusters 4 and 5. In sum, segregation of PVN-CRH cells into several biologically distinct clusters was not driven by ELA-induced changes to transcription.

We then determined if these expression-defined cell subpopulations were differentially impacted by ELA. We computed differential expression between ELA and CTL cells (using an FDR<0.1) separately for each cluster (Supplementary Fig 4A-E). Strikingly, differentially expressed genes following ELA were exclusive to cluster 1. We found 7 genes enriched in ELA cells: *Stmn1, Nnat* (Neuronatin), *Rpl10* (Ribosomal Protein L10), *Vamp2, Hsbp1* (Heat Shock Factor Binding Protein 1), *Sez6l2* (Seizure Related 6 Homolog Like 2), and *Calr* (Calreticulin) in this cluster (Supplementary Fig 4A). While three of these genes were also enriched in the overall population of ELA vs CTL cells, (Figure 1B), four genes (*Nnat, Hsbp1, Sez6l2*, and *Calr*) were specifically enriched in cluster 1 following ELA, demonstrating the increased sensitivity and biological validity provided by the single-cell derived cluster approach. Few genes were enriched in the CTL cells, with a single gene significantly enriched in clusters 1, 2 and 4, cluster 1: *Igf2bp3* (Insulin Like Growth Factor 2 MRNA Binding Protein 3), cluster 2: *Wwc2* (WW And C2 Domain Containing 2), and cluster 4: *Ndufa8* (NADH:Ubiquinone Oxidoreductase Subunit A8) (Supplementary Fig 4A, B, D). Here, the single-cell and cluster approaches identified *Wwc2* and *Ndufa8*, which were not observed in the total-population analyses. Notably, no genes were significantly differentially expressed in clusters 3 and 5 (Supplementary Fig 4C, E).

### Neurotransmitter profiles uncover novel populations of PVN-CRH cells which are selectively vulnerable to ELA

A canonical distinction of neuronal types involves their major neurotransmitter, including glutamate and GABA. The above expression-based clustering suggested that clusters 1, 2 and 4 are primarily glutamatergic, and clusters 2 and 5 GABAergic. Therefore, we determined whether individual CRH+ neurons in developing mouse PVN co-express any or several of these neurotransmitters, and how the neurotransmitter-defined cell populations related to those defined by agnostic gene expression profiles. To this end, we superimposed the Seurat-based clusters onto the expression of glutamatergic markers (the glutamate transporter VGlut2/*Slc17a6*, and the enzyme glutaminase/*Gls*); versus GABAergic markers (the synthesizing enzyme GAD2/*Gad2*; and the vesicular GABA transporter *VGAT/Slc32a1*). As illustrated in Figure 3A, GABAergic CRH+ cells strongly overlapped with clusters 2 and 5, whereas glutamatergic cells dominated the other clusters (*X^2^(4*, N=254) = 52, p < 0.00001). The few cells devoid of any neurotransmitter marker were equally distributed among clusters, suggesting that their expression of neurotransmitter markers simply did not reach detection threshold. Notably, ELA did not significantly change the relative distribution of neurotransmitter-defined subpopulations (*X^2^*(3, N=254) = 4.351, p = 0.226).

**Figure 3.**
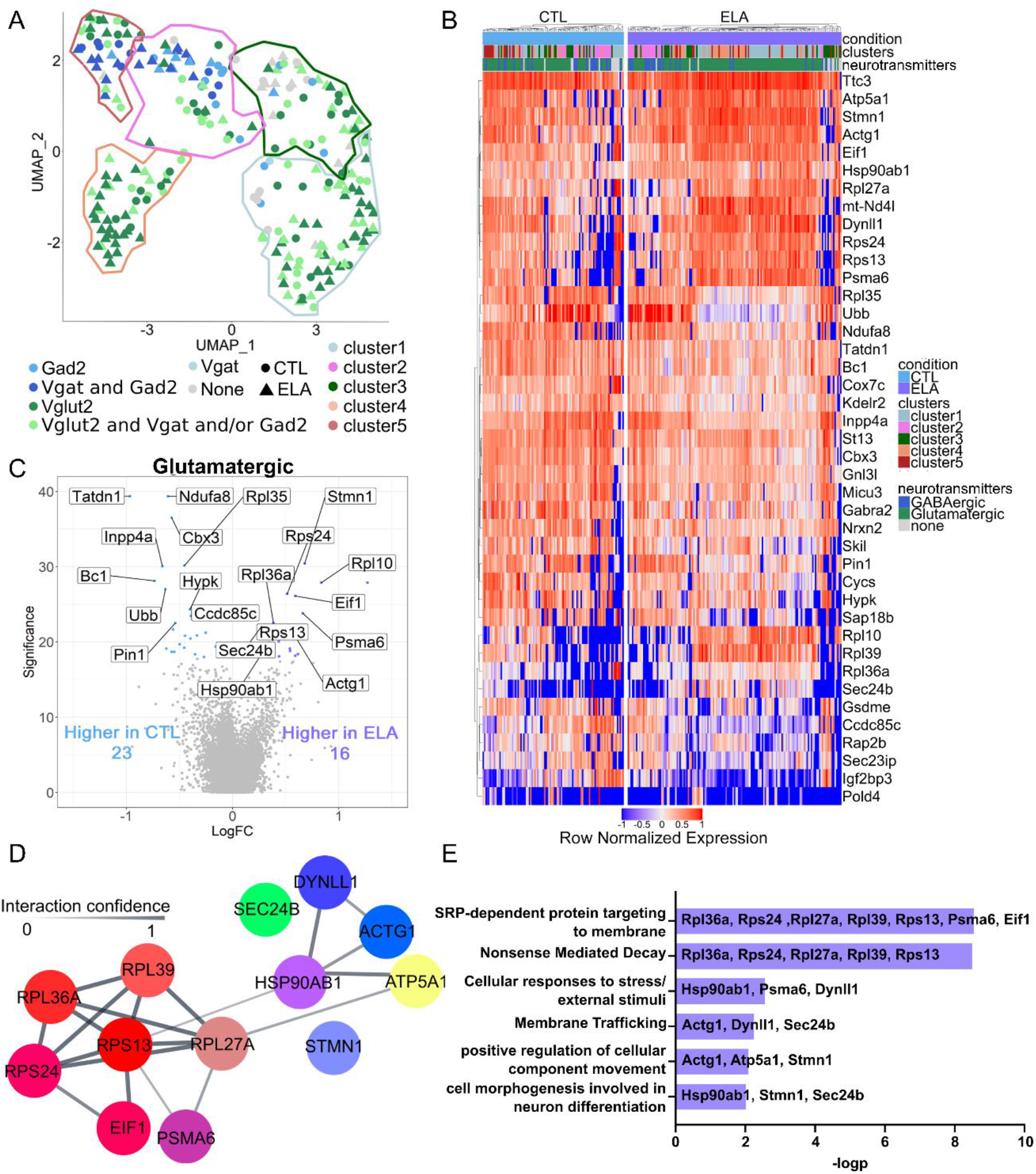
Cell-type specific transcriptomic changes induced by ELA. (A). Expression of neurotransmitters overlayed on the UMAP clustering (Seurat clusters 1-5 circled in colored lines, glutamatergic cells expressing *Slc17a6* and *Gls* in greens and GABAergic cells expressing *Gad2* and *Slc32a1* in blue). (B). Normalized expression of top genes in CTL and ELA cells across the two types of clusters. (C). Genes with significantly increased gene expression changes comparing CTL and ELA glutamatergic cells. (D). Predicted protein-protein interaction (cytoscape) of genes enriched in ELA: nodes represent proteins and edges represent predicted interaction; line strength indicates confidence of predicted interactions from 0-1; cool colors represent genes associated with neuron activation, and warm colors with nonsense-mediated decay (NMD) and protein trafficking. (E). Metascape analysis identifies genes significantly (-logp) associated with predicted pathways which are represented by gene ontology terms.

We then performed differential expression analysis between neurons from the ELA and CTL groups within each neurotransmitter-defined cluster, and found that differentially expressed genes (FDR<0.1) between CTL and ELA cells were highly subtype specific (Figure 3B). There were no differentially expressed genes in the GABAergic cluster, or the cluster comprised of cells expressing no neurotransmitter markers. By contrast, cells within the glutamatergic cluster were transcriptionally modulated by early life experiences. There were 23 differentially expressed genes enriched in CTL cells and 16 enriched in the ELA glutamatergic cells (Figure 3C).

Of the genes enriched in CTL glutamatergic CRH+ cells, 14 were also differentially expressed in the global CTL condition from (Figure 2B), including *Cbx3* (Chromobox 3), *Cox7c* (Cytochrome C Oxidase Subunit 7C), *Gnl3l* (G Protein Nucleolar 3 Like), *Gabra2* (Gamma-Aminobutyric Acid Type A Receptor Subunit Alpha2), *Inpp4a* (Inositol Polyphosphate-4-Phosphatase Type I A), and *Tatdn1* (TatD DNase Domain Containing 1). Genes enriched uniquely in the CTL glutamatergic cluster and not in the global CTL population included *Nrxn2* (Neurexin 2), *Ubb* (Ubiquitin B), *Cycs* (Cytochrome C, Somatic), *Sec23ip* (SEC23 Interacting Protein), *Sap18b* (Sin3A Associated Protein 18), *Pin1* (Peptidylprolyl Cis/Trans Isomerase, NIMA-Interacting 1), and *Ndufa8*. Broadly, the genes upregulated in CTL cells are those involved in the use of the electron transport chain and promote growth: *Cox7c* is required in the oxidative phosphorylation pathway for ATP generation (Timón-Gómez et al. 2018; Ashrafi et al. 2020). *Ndufa8*, the NADH dehydrogenase ubiquinone 1 α subcomplex is heavily involved in mitochondrial oxidative phosphorylation and is expressed in the hypothalamus (Szklarczyk et al. 2011). *Cycs* is the somatic isoform of cytochrome C and is upregulated by neural activity (Delgado & Owens 2012). The differential expression of the ubiquitin encoding gene *Ubb* between CTL and ELA excitatory CRH cells may represent differences in post-translational modifications (Deribe et al. 2010). Gene ontology analysis of these genes upregulated in CTL cells indicated significant enrichment in pathways involved in ATP synthesis-coupled electron transport, positive regulation of intrinsic apoptotic signaling pathway, and regulation of protein binding representing typical cellular metabolism and activity.

Of the 15 differentially expressed genes enriched in the ELA cells, nine were identified in the overall population analyses (Figure 2B; *Actg1, Atp5a1, Eif1, Hsp90ab1, Psma6, Rps13, Rpl10-ps3, Rpl39* and *Stmn1*). Enriched genes unique to the glutamatergic cluster were *Rps24* (Ribosomal Protein S24), *Rpl36a* (Ribosomal Protein L36a), *Sec24b* (SEC24 Homolog B, COPII Coat Complex Component), *Dynll1* (Dynein Light Chain LC8-Type 1), *Nd4l* (Mitochondrially Encoded NADH:Ubiquinone Oxidoreductase Core Subunit 4L), and *Rpl27a* (Ribosomal Protein L27a). Pathway analysis suggested that these genes are associated with response to environmental stressors, regulation of cellular movement and upregulation of translation in ELA cells. The protein interactions (Figure 3D) and functional pathway associations (Figure 3E) for genes upregulated in glutamatergic ELA cells indicate similar gene networks and interaction patterns to those of the total ELA CRH cell population. Namely, in the glutamatergic cell cluster, ELA led to over-representation of genes associated with response to environmental stressors and neuronal activation. *Hsp90ab* was again a central regulator of many of these genes, with predicted associations with gene products involved in protein targeting and nonsense mediated decay as well as cellular response to stress, membrane trafficking, and cellular movement and differentiation (Figure 3 D-E).

### Populations of glutamatergic cells are differentially impacted by ELA

The neurotransmitter-based segregation of CRH+ PVN cells strongly suggested that the glutamatergic cells further belonged to two distinct subgroups (Figure 4A). Therefore, we examined if the impact of ELA on gene expression in glutamatergic cells was selective to one of these sub-populations. The two subclusters of glutamatergic CRH cells were distinguished by their expression of *Avp*, the gene encoding vasopressin, versus *Ntng1*, a cellular adhesion molecule important for axon guidance (Nakashiba et al. 2000; Nakashiba et al. 2002; Zhang et al. 2016), (Figure 4A). Indeed, differential gene expression between ELA and CTL neurons was distinct and non-overlapping in these two glutamatergic cell subsets. In the glutamatergic *Avp* positive sub-cluster (Figure 4B), differentially expressed genes were specifically enriched in CTL cells. These 5 genes included: *Tatdn1*, a long non-coding RNA (lncRNA) important for cell proliferation, adhesion, and migration (Zequn et al. 2016)*; Ndufa8*, which encodes a subunit of NADH dehydrogenase part of the electron transport chain (Szklarczyk et al. 2011); *Cbx3*, a member of the heterochromatin protein 1 family that stimulates cellular differentiation (Huang et al. 2017); the pro-survival gene *Rap2b* (RAP2B, Member Of RAS Oncogene Family) (Qu et al. 2016), and *Bc1* (brain cytoplasmic RNA 1) which is important for translation (Kim et al. 1994). The functions of these genes are required for normal cellular maturation and activity in this subpopulation of CTL cells.

**Figure 4.**
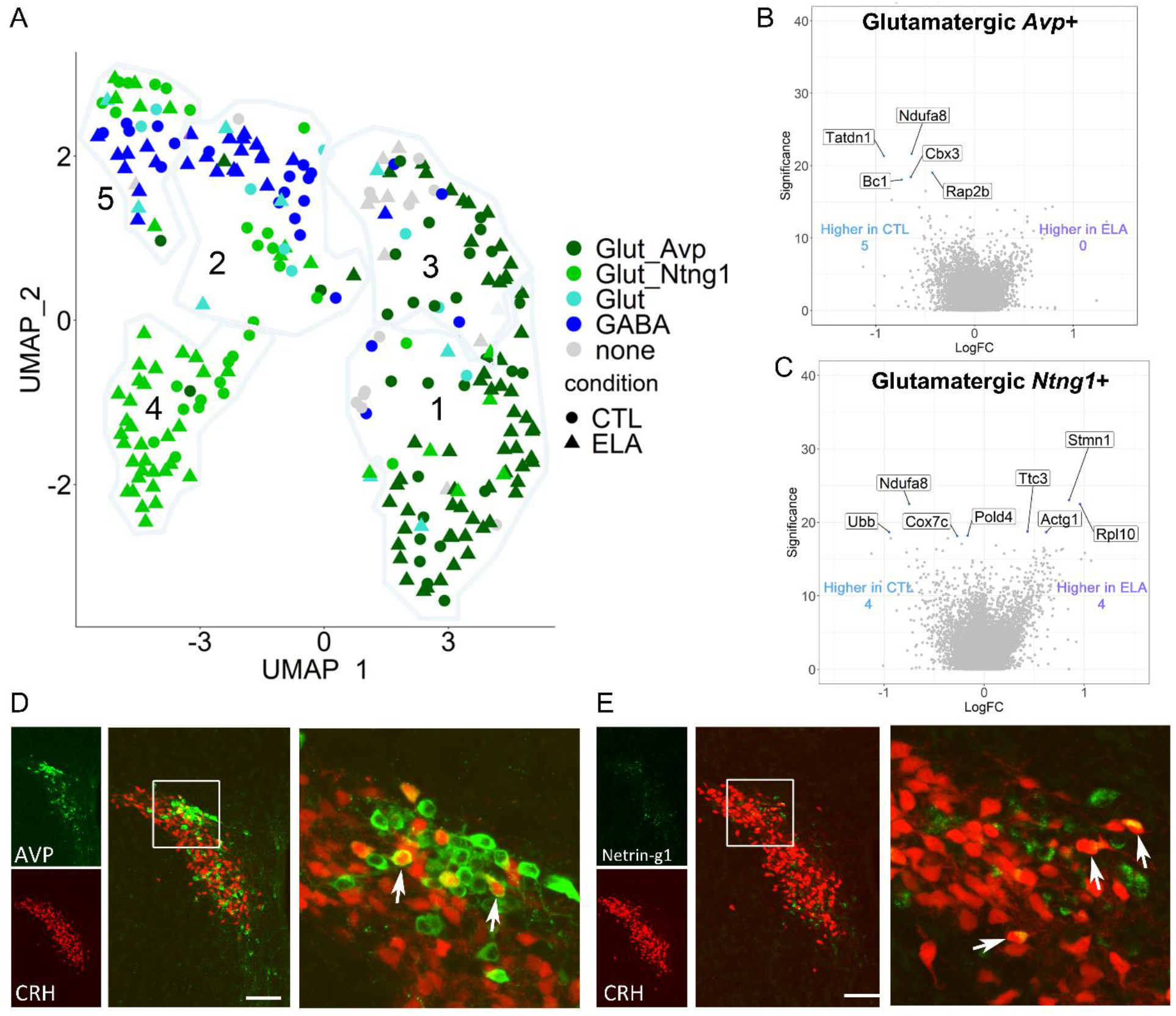
ELA induced transcriptomic changes are specific to novel subpopulations of CRH-expressing glutamatergic neurons, which are differentially spatially organized in the PVN. (A) In CRH expressing PVN cells, neurotransmitter co-expression with *Avp* or *Ntng1* overlayed the UMAP clustering demonstrates distinct glutamatergic subclusters. The Seurat clusters 1-5 are circled in colored lines, glutamatergic CRH+ cells expressing *Slc17a6* and *Gls* in are labeled in greens and GABAergic cells expressing *Gad2* and *Slc32a1* are labeled in blue. (B-C) distinct effects of ELA on gene expression in the AVP and *Ntng1 subclusters:* Genes with significantly increased log fold change in CTL and ELA glutamatergic cells expressing *Avp* (B) and *Ntng1* (C) (CTL=blue and ELA=purple). (D). Dual immunohistochemistry and confocal microscopy demonstrate the spatial organization of neurons co-expressing CRH-promoter driven tdTomato and AVP. (E) pattern of distribution of PVN neurons co-expressing CRH-promoter driven tdTomato and Netring-g1. Co-expressing cells are highlighted by white arrows. Scale bar = 20μM.

**Figure 4.**
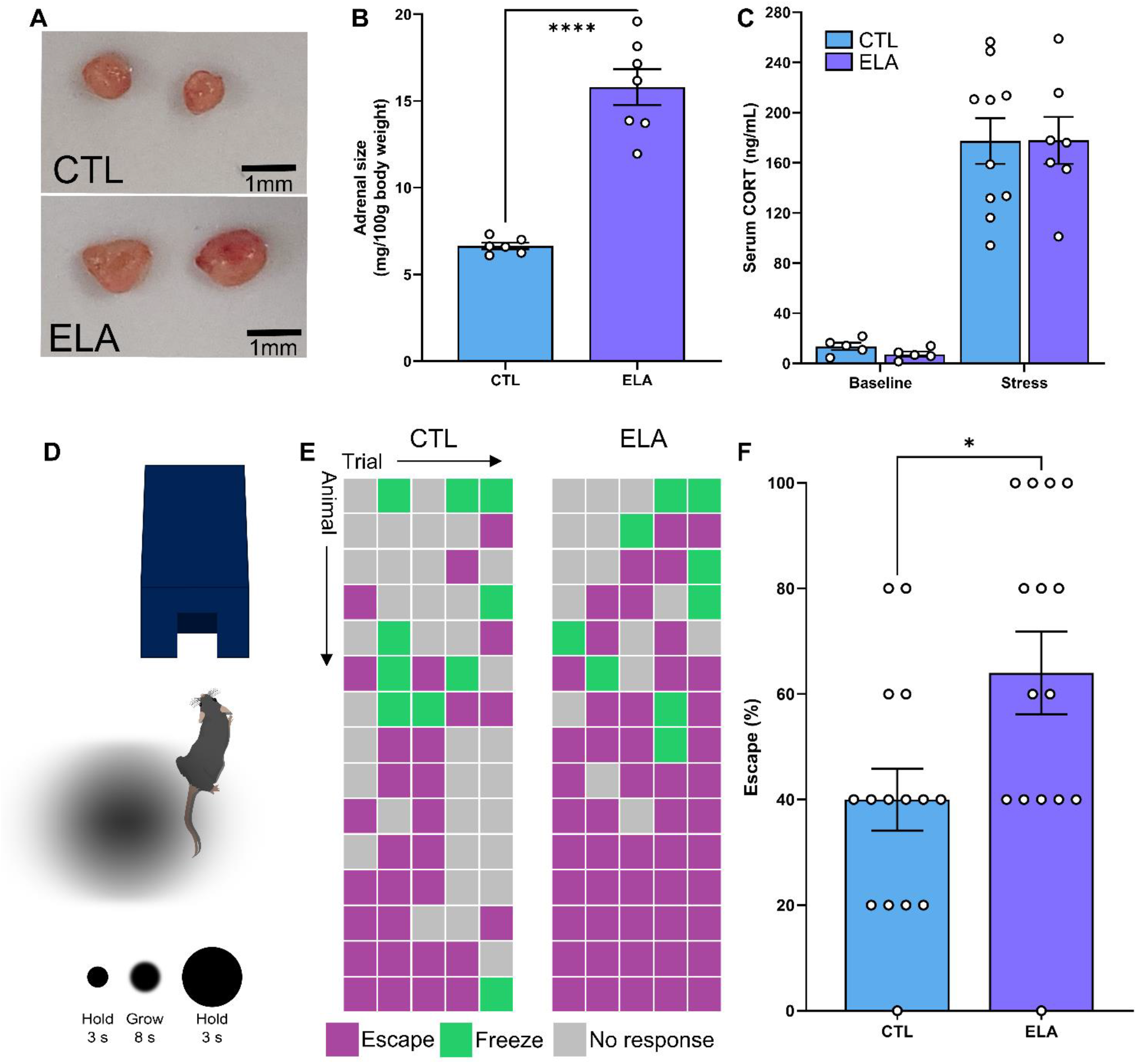
ELA induced Transcriptional changes in CRH expressing PVN neurons herald enduring augmentation of neuroendocrine and behavioral responses to stress. **(A, B)** ELA causes significant increase in adrenal size (n=6-7 per group), a hallmark of a chronic stress state. (C). hormonal responses to acute stress, measured as serum corticosterone at baseline or following one hour of multiple acute stress (MAS), are not influenced by ELA (n=5-10 per group). (D) Schematic of looming disk threat-stimulus test. (E, F) Results from each of the 5-looming threat-stimulus trials for each mouse identify a significant increase in escape behaviors in ELA mice (n=15 per group). Bars represent mean with ±SEM, circles represent individual mice.

Four genes were enriched in the CTL glutamatergic *Ntng1* positive sub-cluster (Figure 4C). These genes included *Ubb, Ndufa8, Cox7c*, and *Pold4* (DNA Polymerase Delta 4, Accessory Subunit), and the latter was uniquely identified in this sub-cluster analysis. *Ndufa8* and *Cox7c* are required for the electron transport train as described above (Szklarczyk et al. 2011; Timón-Gómez et al. 2018; Ashrafi et al. 2020). *Ubb* encodes the highly conserved protein ubiquitin, which controls protein targeting and degradation and has a role in controlling gene expression and the stress response (Finch et al. 1992; Komander 2009). *Pold4* encodes a subunit of DNA polymerase and is critical for DNA replication and repair (Lee et al. 2019). Notably, several genes (*Stmn1, Rpl10*, and *Actg1*) with important neuronal growth functions were uniquely downregulated in the glutamatergic *Ntng1* positive sub-cluster of the controls (enriched in ELA cells; Figure 4C). The robust downregulation of these genes was evident also in the global CRH+ cell population analyses.

### Spatial organization of transcriptionally identified novel populations of CRH+ neurons in the PVN

The discovery of distinct and novel populations of CRH-expressing glutamatergic cells in the PVN, which are differentially influenced by ELA, relied on single-cell transcriptomics and cluster analyses. Therefore, we sought to determine the physical distribution of these populations, focusing on colocalization of AVP and of Netrin-g1 with CRH. To this end, we performed immunohistochemistry on slices from mice during the developmental period of the sequenced cells. As shown in Figure 4D, a subset of PVN neurons expressing tdTomato under the CRH promoter co-expressed AVP. These CRH-AVP neurons resided primarily in the dorsal PVN. By contrast, cells expressing CRH together with Netrin-g1 were distributed sparsely throughout the PVN, without apparent spatial organization (Figure 4E).

### Transcriptional changes provoked by ELA herald enduring significant alterations of behavioral and hormonal responses to stress

To determine the potential functional long-term significance of the transcriptional changes induced by ELA in glutamatergic PVN-CRH cells, we determined behavioral and hormonal responses to stress in ELA and CTL mice later in life. We measured chronic increase of the release of CRH and its downstream stress hormones using adrenal weights (Ulrich-lai et al., 2006) as well as hormonal responses to acute stress. Adrenal gland weights of adult mice that experienced ELA were significantly higher than those of control mice (t_(11)_=8.1, p<0.0001), (Figure 5A), in accord with our prior work (Avishai-Eliner et al. 2001; Brunson 2005), suggesting chronically heightened response to stress throughout life (Ulrich-Lai et al. 2006). Responses to acute stress were determined using exposure to multiple acute concurrent stresses imposed for one hour (Chen et al. 2016; Hokenson et al. 2020), which induce robust corticosterone secretion in adult male mice (Maras et al. 2014). (Figure 5B). As expected, serum corticosterone levels were higher following acute stress than at baseline (F_(1,23)_=89.11, p<0.0001), yet there was no main effect of early-life adversity on serum corticosterone (F_(1,23)_=0.03, p=0.87), and no stress by ELA interaction (F_(1,23)_=0.03, p=0.85) (Figure 5B). Together these experiments indicated that transcriptional changes detectable already immediately after transient ELA result in enduring and robust changes in the chronic but not acute neuroendocrine responses to subsequent stresses. Whereas responses to severe acute stress do not distinguish adult ELA from control mice, the ELA mice release much more hypothalamic CRH and pituitary ACTH chronically, resulting in adrenal hypertrophy, the hallmark of a chronic stress state.

The functional role of stress-sensitive PVN-CRH cells in orchestrating behavioral responses to stress has not been fully elucidated. A requirement of these neurons d in behavioral responses to threat such as fleeing or freezing was recently shown (Daviu et al. 2020b). Therefore, tested the consequences of ELA on behaviors mediated by PVN-CRH cells by employing a test for eliciting threat responses, the looming shadow task (Daviu et al. 2020a). In this test, escape behaviors of male mice raised under ELA conditions were significantly higher compared to controls (t_(22)_=2.449, p=0.02) (Figure 5C-D). These data suggest that the transcriptional changes resulting from ELA may contribute to altered behavioral response to stressful threats later in life.

## Discussion

There are several principal findings of the current studies, which employ single-cell transcriptomics analysis of CRH expressing PVN neurons and determine the impact of adverse / stressful early life on gene expression patterns. We found that CRH-expressing neurons in the PVN can be clustered into distinct populations by both their gene expression profiles as well as by their neurotransmitter phenotype, and these populations are distinctly spatially organized in the PVN. ELA modified gene expression patterns, impacting transcriptional programs of neuronal development and differentiation and enhancing gene families involved in responses to stress and inflammation. The use of single-cell transcriptomics revealed that ELA impacts gene expression profiles in a cell-type specific manner, with distinctive influence on the different clusters and subpopulations of CRH neurons. Finally, the transcriptional changes identified immediately following ELA heralded significant life-long disruptions of the hormonal and behavioral responses to stress.

### PVN contains discrete, molecularly defined clusters of CRH cells

Clustering using Seurat identified differences in gene expression profiles of CRH positive PVN neurons, which may be consistent with those previously described (Romanov et al. 2017a; Chen et al. 2017; Xu et al. 2020; Romanov et al. 2020). Clusters 1 and 3 had increased expression of *Avp* and steroid hormone receptors. Whereas the transcriptomic profile of cluster 5 would suggest these represent a small population of cells which co-express GABA and CRH within the PVN. Previous studies have found that while the molecular defined clusters were not representative of spatial location within the PVN, some of the observed transcriptomic changes may be indicative of cells performing different functions or that are sampled at differing stages of development (Simmons & Swanson 2009; Xu et al. 2020).

Specifically, clusters 1 and 3 may include populations of cells undergoing maturation and differentiation. Cluster 1 was highly enriched in genes encoding ribosomal proteins required for protein targeting and translation (Nosrati et al. 2014). Cluster 3, the second cluster with high expression of *Avp*, included differentially expressed genes associated with cell proliferation, differentiation, and activation. Expression of genes in this cluster include the immediate early gene *Junb* (JunB Proto-Oncogene, AP-1 Transcription Factor Subunit), which is essential for neuronal differentiation (Schlingensiepen et al. 1994) and the neuropeptide *Npff*, which decreases synaptic drive to magnocellular AVP-secreting PVN neurons (Jhamandas & Goncharuk 2013). Cluster 3 also had enrichment of *Dlk1* (delta like non-canonical Notch ligand 1), which is expressed in AVP and oxytocin secreting PVN neurons in both adult and postnatal mice and has been associated with neuroendocrine differentiation (Villanueva et al. 2012).

Cluster 4 had an over-representation of cells from the ELA group. Following ELA, there is a significant increase in the formation of excitatory synapses on the CRH cells of the PVN (Gunn et al. 2013; Bolton et al. 2021). Many of the genes upregulated in cluster 4 could indicate increased synapse development. For example, transcription factors such as *Tcf7l2* (Transcription Factor 7 Like 2), *Zic2* (Zinc Finger Protein Of The Cerebellum 2) *and* Prox1 (Prospero Homeobox 1) that are involved in neuronal identity and circuitry, axonal navigation, neuronal migration, and neurogenesis (Kaltezioti et al. 2010; Murillo et al. 2015; Lee et al. 2017), were enriched in cluster 4. *Ntng1* was also highly expressed in this cluster, similar to clusters observed in previous studies (Xu et al. 2020). The encoded protein, netrin-G1, is a synaptic cell adhesion molecule that is enriched in presynaptic terminals (Kim et al. 2006; Zhang et al. 2016). Like the netrin family, netrin-G1 is an axon guidance molecule, however, it does not interact with traditional netrin receptors (Lin et al. 2003; Kim et al. 2006). The binding of netrin-G1 with its ligand NGL1 is important in growth of embryonic thalamic axons (Lin et al. 2003), this could suggest that cluster 4 represents cells actively forming synapses. The observed increase above chance of ELA cells within this cluster could represent the increased formation of synapses in CRH neurons following ELA.

Cluster 5 genes were representative of axon development, microtubule development, and synaptic transmission. Genes enriched in this cluster include the tubulin family of genes, including *Tubb2a* (Tubulin Beta 2A Class IIa) and *Tubb2b* (Tubulin Beta 2B Class IIb), these are required for microtubule development, which is essential for many cellular processes, especially during neural development (Bittermann et al. 2019). *Vamp1* (Vesicle Associated Membrane Protein 1) is involved in the presynaptic docking of vesicles and is required for Ca2+-triggered synaptic transmission (Liu et al. 2011). The genes expressed in this cluster may represent neurons undergoing increased activity.

### Adversity during sensitive periods enriches genes associated with neuronal maturation, differentiation, and stress

Analysis of the entire CRH cell population revealed specific changes in gene expression profiles between the ELA and the CTL condition. Our results suggest that while control cells are engaged in typical cellular functions, cells from the ELA condition are enriched for genes important for neuronal maturation, differentiation, and mitigating neuronal stress. Genes such as *Cox7c* and *Micu3* that are required in the oxidative phosphorylation pathway for ATP generation (Timón-Gómez et al. 2018; Ashrafi et al. 2020), were enriched in CTL cells. *Cbx3*, a gene important for neural differentiation, was also significantly upregulated in CTL cells. *Cbx3* increases expression of neuronal genes and inhibits expression of genes specific to other fates (Huang et al. 2017). This upregulation of *Cbx3* in CTL cells and the concurrent downregulation in ELA cells, suggests a potential neuronal de-differentiation of CRH cells in the PVN consequent to ELA, similar to the observation in hippocampus (Bolton et al. 2021). A neuronal fate is highly ‘expensive’ metabolically, requiring energy-intensive maintenance of membrane potential and neuronal firing. De-differentiation to another cell fate may save the cell from death in conditions of stress.

Contrary to the transcriptomic profile representing typical cellular functions observed in control cells, cells collected from animals following ELA have profiles consistent with response to stress and neuronal activation (Figure 2 D-E). The HSP90 family of heat shock proteins are important for the facilitation of steroid hormone receptor function. HSP90 maintains the glucocorticoid receptor (GR) in the high affinity binding state and is required for GR translocation to the nucleus (Pratt et al. 2006; Noguchi et al. 2010). The upregulation of *Hsp90ab1* in ELA animals may make it an important regulator in the signaling cascades responsible for the cellular activation following stress and subsequent protein changes.

### ELA upregulated genes associate with increased activity in glutamatergic excitatory CRH cells

The clustering analysis presented here highlights the complexity of CRH expressing cells within the PVN (Figure 1E), and that the clusters characterized here identify neuronal populations beyond those described previously. Using the highly expressed genes identified for each cluster, the cells were subdivided based on their neurotransmitter expression patterns (Figure 3A). As described previously (Ziegler et al. 2005; Wamsteeker Cusulin et al. 2013; Chen et al. 2015; Füzesi et al. 2016), PVN CRH-expressing cells were predominately glutamatergic, and included also a small population of cells expressing GABAergic markers. Remarkably, the effects of ELA were only observed in the population of glutamatergic cells.

What might the ELA-induced changes in gene expression patterns of glutamateric CRH cells signify? Following ELA, there is a significant increase in functional excitatory synapses on the CRH cells of the PVN (Gunn et al. 2013; Bolton et al. 2021). This increase should drive both increased metabolic stress within these cells as well as potentially increase in their response to input from other brain regions that process stress throughout life. The changes provoked by ELA support both of these possibilities: First, the differentially expressed genes enriched in the ELA condition uniquely in the glutamatergic cluster (*Dynll1, Nd4l, Rpl27a, Rpl36a, Rsp24*, and *Sec24b*) largely encode proteins required for transcription and translation complexes (Bhavsar et al. 2010; van Oterendorp et al. 2011; Baines et al. 2013). Gene ontology identified nonsense mediated decay (NMD) as a pathway significantly upregulated in the glutamatergic CRH cells following ELA. NMD regulates the expression of proteins related to targeted degradation of mRNA in conditions of cell stress and apoptosis. NMD has also been implicated in regulation of synaptic plasticity (Notaras et al. 2020).

Further, among the two subclusters of glutamatergic cells that were defined based on expression of *Avp* or *Ntng1* (Figure 3) all of the ELA-induced gene expression enrichment took place in the *Ntng1* expressing cells. Thus, the *Ntng1* positive cells are driving the transcriptomic changes observed following ELA. Netrin-G proteins are important for regulating and diversifying synapses. Loss of netrin-G1 - NGL1 interaction causes decreased synaptic plasticity of excitatory inputs in hippocampus (Matsukawa et al. 2014). Additionally, mice lacking netrin-G1 exhibit a reduction in anxiety-like freezing responses in cued and contextual tests, and impairments in goal directed strategy on the Morris water maze with no deficit in spatial learning (Zhang et al. 2016). Although the function of the *Ntng1* positive cluster of CRH cells within the PVN is unknown, the fact that ELA-provoked changes target cells rich in netrin-G1, a regulator of synapse development, is congruent with—and may mediate--the aberrant increase in excitatory synapses onto ELA CRH+ cells. The functional changes described here also indicate an increase in activation of the PVN CRH cells. Increased synapses on CRH cells within the PVN would lead to increased activity of these cells in response to everyday stressors. Over the lifespan, this heightened stress response may result in the increased adrenal size, often associated with chronic stress (Ulrich-Lai et al. 2006), observed here. The lack of observable difference in serum corticosterone in this study may represent compensatory mechanisms or a lack of sensitivity in collection time. The increased propensity for ELA animals to escape during the threat response test also suggests increased activity of the PVN CRH cells. Previous work has linked the activation of PVN CRH cells with escape behavior in the same looming shadow test. Specifically, optogenetic inhibition of PVN-CRH cells attenuated escape behaviors (Daviu et al. 2020b). In contrast, in the current study ELA resulted in significantly more escape responses compared to control mice.

In conclusion, the use of single-cell transcriptomic enables an unprecedented level of insight into the diverse CRH+ expressing neuronal populations within the PVN, as well as the altered gene expression patterns provoked by ELA in these neurons. These involve cellular processes such as response to stress, synapse formation and altered energy production, and may reflect the abnormal increased number of excitatory synapses observed neuroanatomically in these cells after ELA. This work highlights a new level of heterogeneity of the CRH cells within PVN and segregates these cell population into biologically meaningful clusters with distinct spatial organization. Indeed, several of these cell subsets are particularly sensitive to adversity early in life. Understanding which cell types undergo transcriptional programming in response to early environmental signals, and how these experiences are encoded transcriptionally is vital for identifying novel interventions to mitigate the enduring effects on brain development.

## Methods

### Animals

Dams were Crh-IRES-Cre +/+ (Taniguchi et al. 2011) and were paired with Ai14 tdTomato (Madisen et al. 2009) males both on a C57Bl6 background. The resulting offspring were Crh-IRES-Cre; Ai14 tdTomato, which express tdTomato with nearly full overlap of native CRH (Wamsteeker Cusulin et al. 2013; Chen et al. 2015), and were used for all experiments. Animals were housed in a 12-hour light cycle and provided *ad libitum* food and water. All experiments were carried out in accordance with the Institutional Animal Care and Use Committee at the University of California-Irvine and were consistent with Federal guidelines.

### Early-life adversity paradigm-cages with limited bedding and nesting (LBN)

**We imposed ELA on neonatal mice using** simulated poverty by limiting nesting and bedding materials in cages between P2-P9 as described previously (Gilles et al. 1996; Rice et al. 2008; Molet et al. 2014). For the LBN group, a plastic-coated mesh platform was placed ~2.5cm above the floor of a standard cage. Cobb bedding was reduced to cover the cage floor sparsely, and one-half of a single nestlet was provided for nesting material on the platform. Control dams and litters resided in standard cages containing ample cob bedding and one whole nestlet for nesting. This paradigm causes maternal care to be fragmented and unpredictable (refs), provoking chronic stress in the pups (refs). Control and experimental cages were undisturbed during P2–P9, housed in temperature-controlled rooms (22°C). For RNA sequencing pups remained on the LBN paradigm until tissue was collected P10-P12. For experiments in adulthood, experimental groups were transferred to standard cages on P10 and were weaned on P21. Animals were housed by sex with littermates.

### Single cell preparation

#### Dissection

P10-P12 pups were killed via decapitation and brains were removed immediately on ice. The brain was trimmed to a smaller block containing hypothalamus and was placed into a slush of EBSS (*NaCl 116 mmolL^-1^, KCl 5.4 mmolL^-1^, CaCl_2_ 1.8 mmolL^-1^, MgSO_4_.7H_2_O 0.4 mmolL^-1^, NaHCO_3_ 26.2 mmolL^-1^, Glucose 5.5 mmolL^-1^).* The block was then sliced on a vibratome to obtain at 1.5mm slice posterior of the anterior commissure. The slice was then trimmed under a dissection microscope to remove thalamus and cortex.

#### Dissociation

The trimmed slices were placed immediately into papain (add strength) and gently triturated ~15 times with a pipette to break up tissue before being placed on a heated (37C) orbital shaker for 30 minutes. Digested tissue was then homogenized by triturating with pipette ~ 50 times until there were no visible tissue chunks. Homogenate was spun at 500 x g for 15 minutes at room temperature. Supernatant was removed and cells were resuspended in 500uL 2% FBS in PBS.

#### FACS

Samples were run on the BD FACSaria fusion (BD Biosciences, NJ USA). Immediately prior to sort, cell suspension was run through a 70uM filter and washed with 500uL 2% FBS in PBS. ADD SORT DETAILS. Cells were sorted into 8 well strip tubes and immediately spun down at 4C and frozen on dry ice.

### RNAseq pipeline

Tdtomato positive hypothalamic cells from both ELA and CTL animals were processed using the SmartSeq2 RNA sequencing protocol, Illumina Library prep and sequenced using the Illumina NextSeq500 sequencer (Illumina, San Diego) to an average depth of 3.8 million reads per cell. Reads were mapped and quantified using kallisto (Bray et al. 2016). Cells were filtered for a minimum expression of 1000 genes per cell, and only genes expressed in 4 or more cells were included. The resulting TPM matrix was quantile normalized before using the R package Seurat to cluster and run non-linear dimensional reduction (Stuart et al. 2019). ComplexHeatmap was used for heatmap generation (Gu et al. 2016). Metascape (Zhou et al. 2019) was used to determine gene ontology and pathways.

### Immunofluorescence staining

P10 pups were euthanized with sodium pentobarbital and transcardially perfused with ice-cold phosphate-buffered saline (PBS; pH=7.4) followed by 4% paraformaldehyde in 0.1M sodium phosphate buffer (pH=7.4). Perfused brains were post-fixed in 4% paraformaldehyde in 0.1 M PBS (pH = 7.4) for 4-6 hr, before cryoprotection in a 25% sucrose solution. Brains were frozen, then sectioned coronally into 20-μm-thick slices (1:6 series of the PVN) using a Leica CM1900 cryostat (Leica Microsystems, Wetzlar, Germany). Immunofluorescence staining was performed on brain sections derived from P10 male tdTomato-Crh (*Crh-IRES-Cre;Ai14*) transgenic mice as described previously (Chen et al. 2015; Gunn et al. 2019). Briefly, after several washes with PBS containing 0.3% Triton X-100 (PBS-T, pH 7.4), sections (20 μm) were treated with 0.3% H2O2/PBS for 30 min, then blocked with 5% normal goat serum (NGS) for 30 min to prevent non-specific binding. After rinsing, sections were incubated for 3 days at 4°C with rabbit anti-AVP (1:2,000, Sigma, PC234L) or rabbit anti-Ntng1 (1:1,000, ThermFisher, PA5-30447) in PBS containing 1% BSA, and washed in PBS-T (3 × 5 min). Immunoreactivity was visualized using anti-rabbit IgG conjugated to Alexa Fluor 488 (1:400, Invitrogen) for 2 hrs (RT).

### Confocal imaging

Brain sections from the PVN (equal to AP −0.46 mm to −0.94 mm in the adult, relative to the Bregma) were subjected to confocal imaging (LSM 510, Zeiss). Virtual z-sections of 1 μm were taken with an Apochromat 63× oil objective (numeric aperture 1.40). Image frame was digitized at 12 bit using a 1024 x 1024 pixel frame size. To prevent bleed-through in dual-labeling sections, images were scanned sequentially by two separate excitation laser beams: a Argon laser at a wavelength of 488 nm and a He/Ne laser at 543 nm. Z-stack reconstructions and adjustments of image contrast were performed using ImageJ (version 1.41).

### Tests of CRH+PVN cell function

#### Looming shadow task

The looming shadow task is largely dependent on the CRH neurons of the paraventricular nucleus (PVN) of the hypothalamus (Daviu et al. 2020b). The task was performed as described previously (Daviu et al. 2020a). Briefly, animals were initially habituated to an arena containing a shelter for 15 minutes during their dark cycle. Following habituation, a stimulus is presented to the animal from above which consisted of an expanding disk (add exact times) which was repeated 5 times with at least 1 minute between stimuli. The animal’s response to the stimulus is then scored (eg. no response, freezing, escape) both live and on recording by an independent experimenter.

#### Stress in adulthood

Responses to acute stress were determined using multiple acute concurrent stresses (Chen et al. 2016; Hokenson et al. 2020) imposed for one hour. This paradigm involves exposing mice to simultaneous physical, emotional, and social stresses and is described in detail at Bio-protocol (Hokenson et al. 2020) and has been utilized in other studies (Maras et al. 2014; Chen et al. 2016; Hokenson et al. 2019; Libovner et al. 2020). Briefly, mice were individually restrained in a ventilated 50mL plastic tube. Two to six mice were placed in a cage atop a laboratory shaker in a room with loud (90dB) rap music and bright lights for one hour. Animals were perfused immediately post-stress.

#### Corticosterone assay

Blood was collected through cardiac puncture prior to perfusion and samples were left to clot at room temperature for 30 minutes. Following centrifugation at 1100 x g for 15 minutes serum was collected at stored at −20°C. Corticosterone in the serum was analyzed using the Corticosterone EIA Kit (Cayman Chemical Company, MI, USA) according to the manufacturer’s instructions. Post-stress serum was collected directly following the stress. Optimal serum dilutions were previously established in the lab and a 1 in 50 dilution for basal levels and 1 in 500 for post-stress serum was analyzed in duplicate.

#### Adrenal gland collection

In a separate cohort of mice, adrenals were collected following a lethal injection (Euthasol solution; ~488 mg/kg pentobarbital sodium and ~63 mg/kg 175 phenytoin sodium, intra-peritoneally). Gross dissection was performed to isolate the left and right adrenals from the surrounding tissue and were weighed together. Adrenal size is expressed as a function of body weight collected at the time of euthanasia.

### Statistical considerations and analyses

Where possible, data collection and analyses were performed blind to treatment group. Statistical analyses for chi squared tests and functional tests were performed using GraphPad prism 9.0 (GraphPad software, Inc., LA Jolla, CA), using a student’s t-test with significance set at p≤0.05. All other analysis were performed using R studio version 4.0.2 (RStudio Team 2020). Graphs show the mean ± standard error of the mean (SEM).

## Supplementary Figures

**Supplementary Figure 1.**
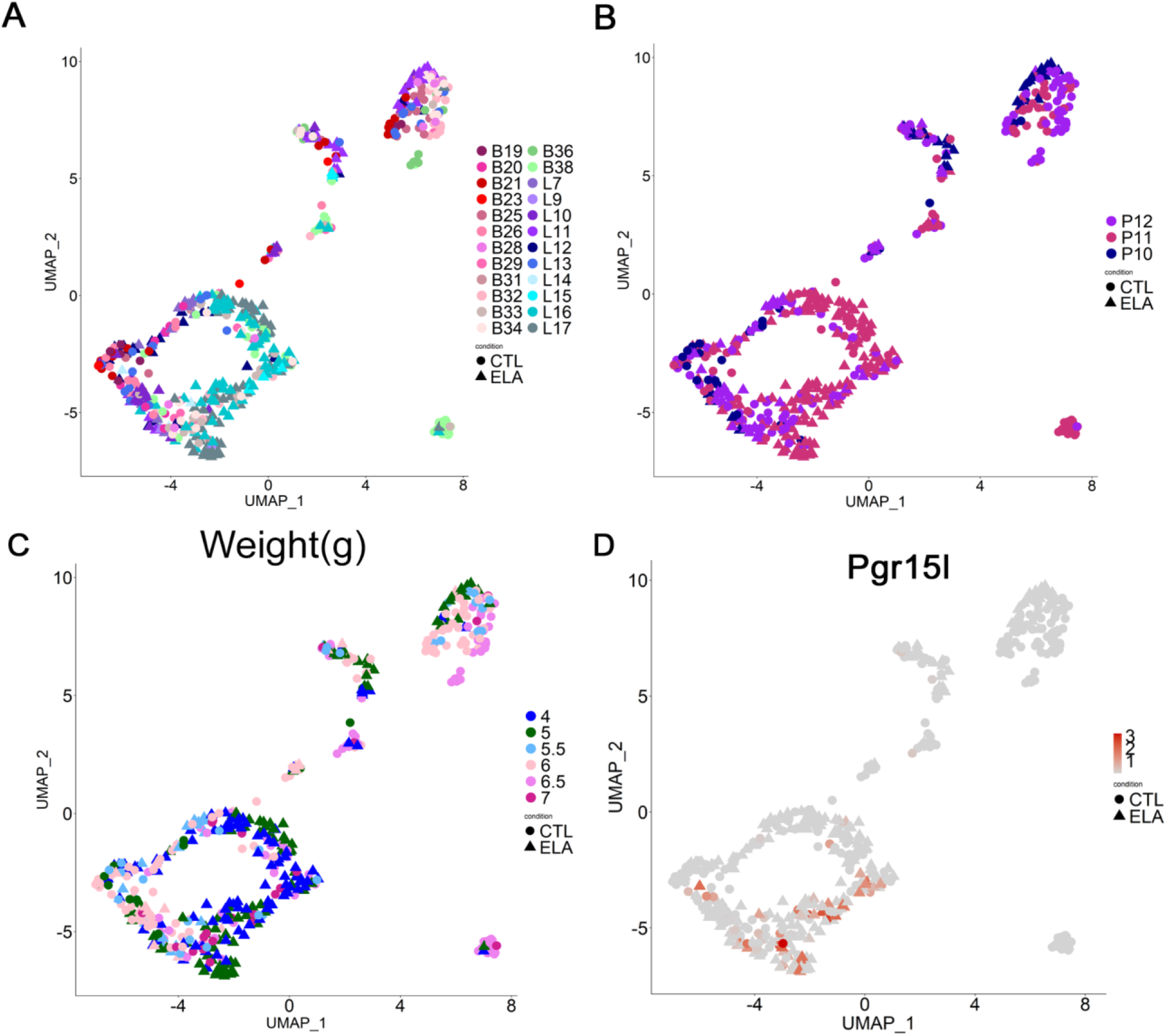
Characterization of batch, age, weight, and non-PVN marker *Pgr15l* expression of 511 cells. A) UMAP characterization of 24 batches across 511 cells to determine little or no batch effect between sorts. B) UMAP characterization of age the cells were harvested between postnatal day 10 - 12. C) UMAP characterization of weight in grams of mice on the day cells were harvested. D) Heatmap overlay of non-PVN *Pgr15l* expression over the UMAP of 511 cells to determine cell filter for *Pgr15l* expression. All UMAPs were generated using Seurat v3.

**Figure 2.**
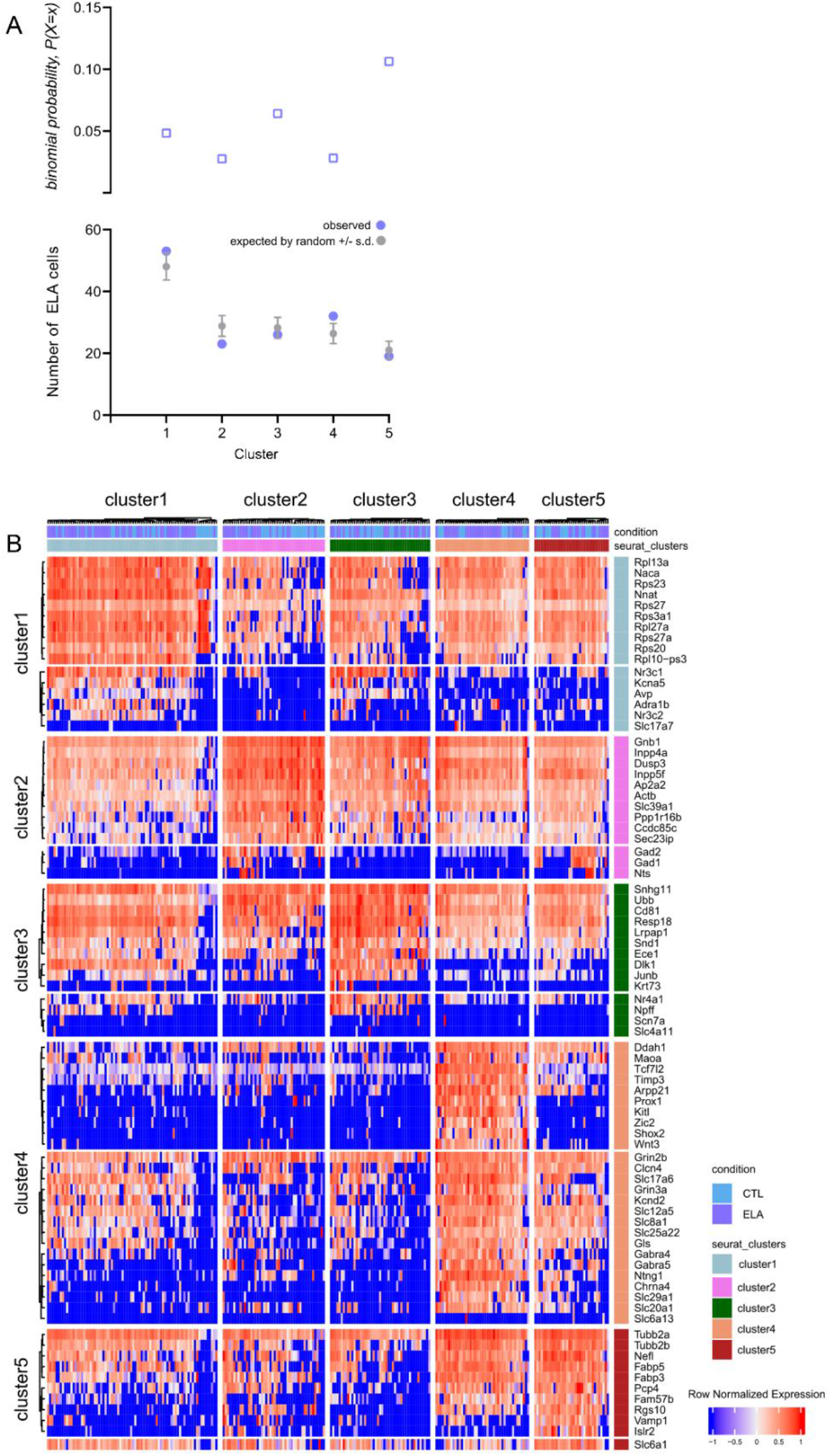
Distribution of cells and genes between clusters. (A) the observed number of ELA cells in each cluster compared to what would be expected at random according to binomial distribution and the probability of this occurring by chance. (B) Top 10 genes enriched in each cluster (top panels for each cluster) and identified genes of interest (bottom panels for each cluster), row log normalized.

**Figure 3.**
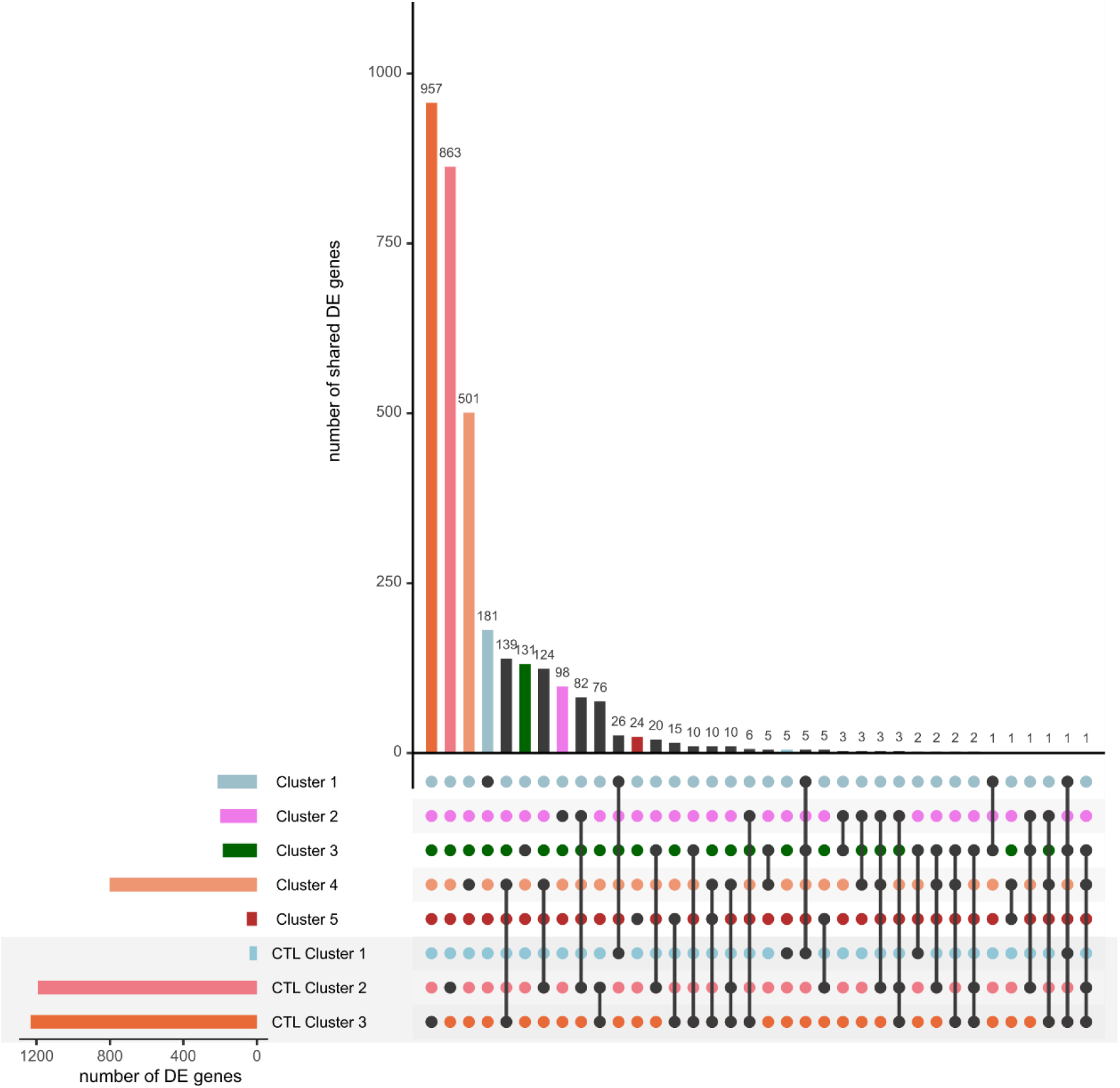
Representation of the number of shared differentially expressed genes per cluster. Most clusters including CTL and ELA cells had distinct DE genes; clusters only including CTL cells (in the grey box) largely encompassed the genes from the ELA clusters.

**Figure 4.**
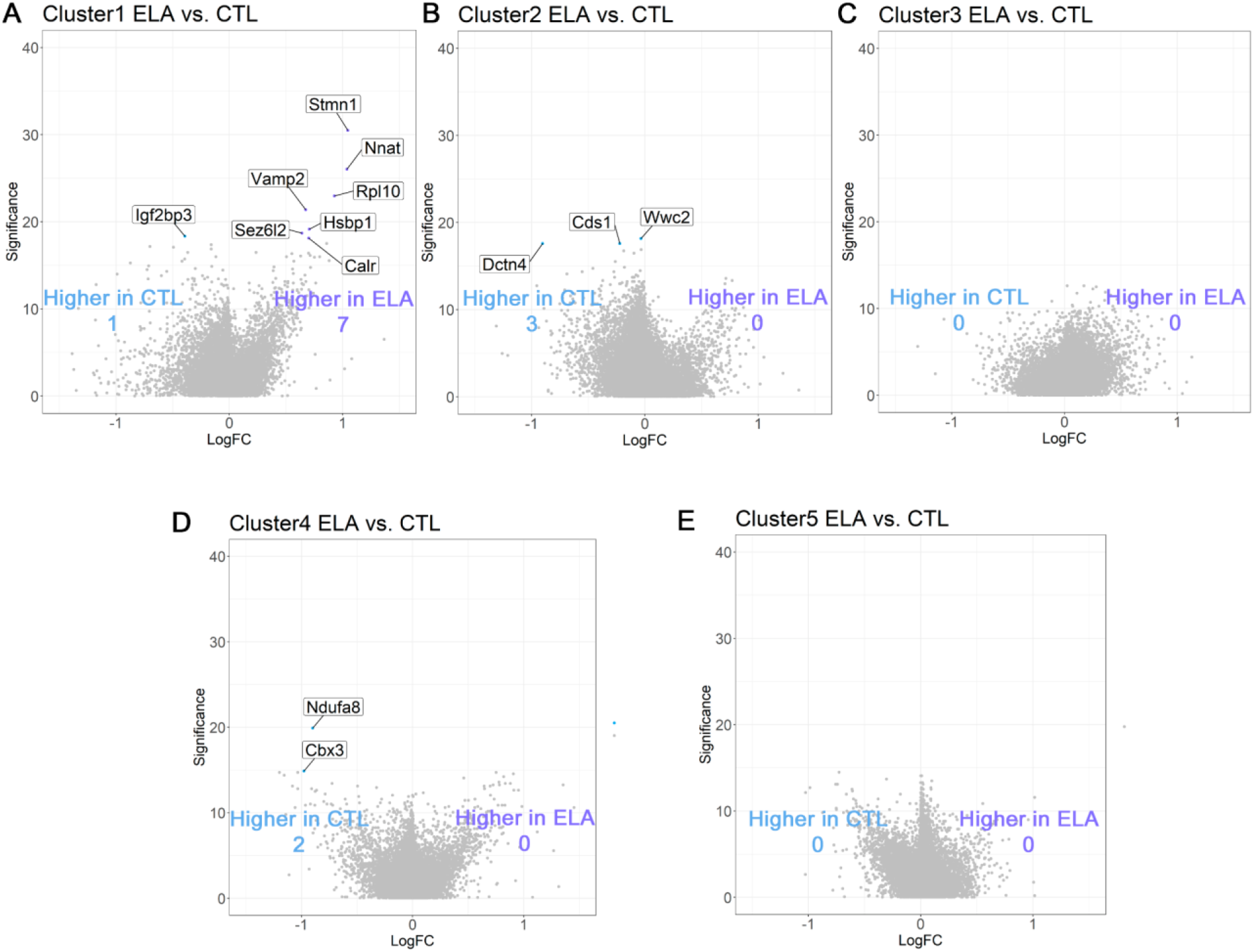
Differential expression analysis in each Seurat cluster between the ELA and CTL cells. Volcano plots representing differential expression between the ELA and CTL cells. (A) In cluster 1, there were 7 differentially expressed genes enriched in the ELA cells and 1 in the CTL cells. B) In cluster 2, there were 0 differentially expressed genes enriched in ELA cells and 3 in the CTL cells. C) No differentially expressed genes were observed in cluster 3. D) There were 0 differentially expressed genes enriched in ELA cells and 2 in the CTL cells in cluster 4. E) No differentially expressed genes were present in cluster 5. All enriched genes were assessed for false discovery rates (FDR) <0.1.

